# Human iPSCs from aged donors retain their mitochondrial aging signature

**DOI:** 10.1101/2024.04.16.589733

**Authors:** Imane Lejri, Zameel Cader, Amandine Grimm, Anne Eckert

## Abstract

Aging represents the main risk factor for developing neurodegenerative disorders. One of the hallmarks of aging is mitochondrial dysfunction. Age-related mitochondrial alterations have been shown to affect mitochondrial energy metabolism and redox homeostasis as well as mitochondrial dynamics. In the present study, we addressed the question of whether or not, induced pluripotent stem cells (iPSCs) may be used as a model of “aging in a dish” to identify therapies at alleviating the aging of mitochondria. Notably, we could demonstrate that compared to human iPSCs from young donors, those from aged donors show impaired mitochondrial bioenergetics and exhibit a rise in reactive oxygen species generation. Furthermore, we demonstrate that iPSCs from aged donors present low mitochondrial mass and alterations of the morphology of the mitochondrial network. This study provides evidence that the aging phenotype is present at the mitochondrial level in iPSCs from aged donors, ranging from bioenergetics to dynamics. Thus, this model can be used for high through put screening to identify drugs that improve mitochondrial function.

**Graphical abstract:** 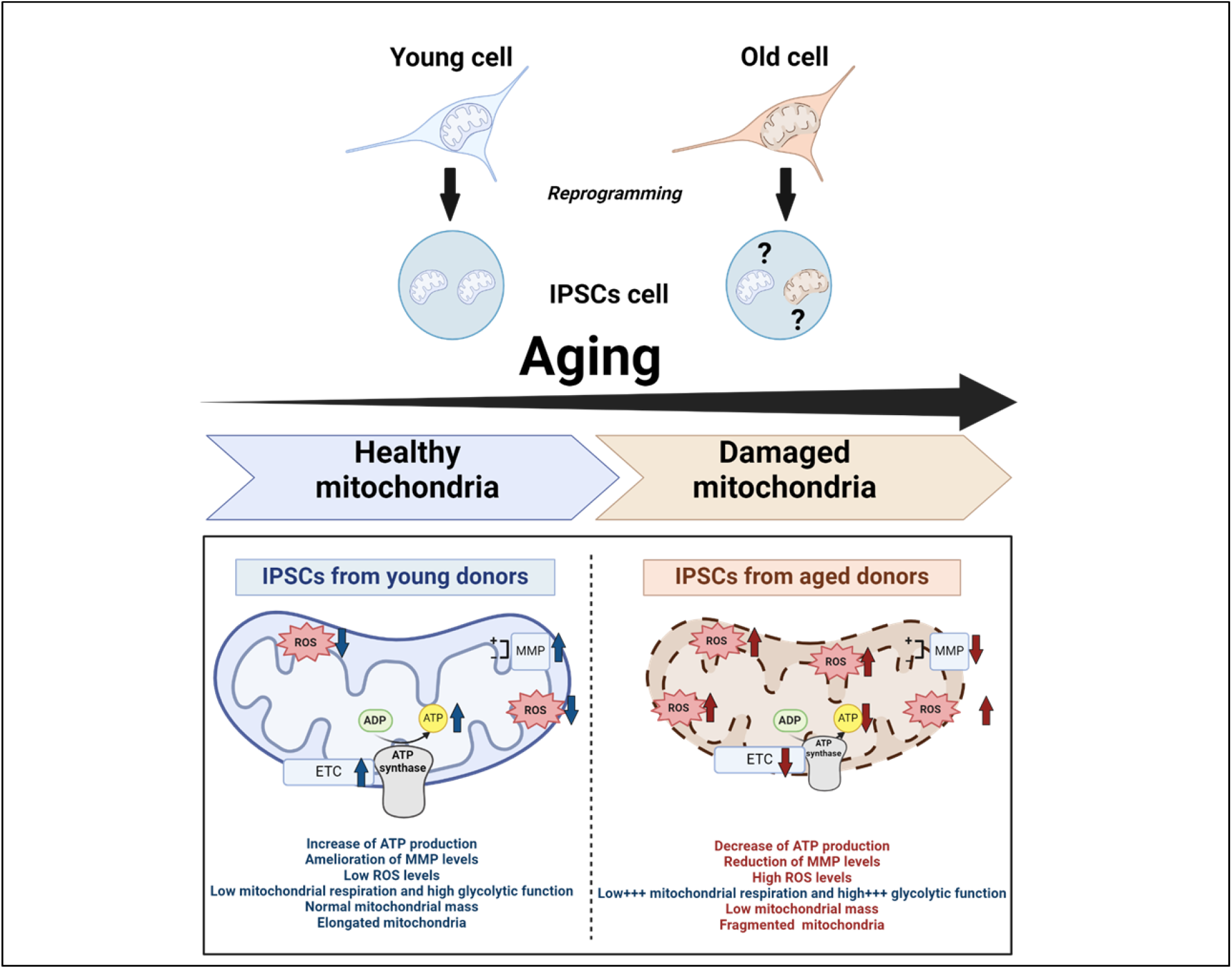

## 1. Introduction

In aging, several hallmarks are reported including the mitochondrial dysfunction that is also known to be related with the evolution of age-related neurodegenerative disorders (Kauppila, Kauppila et al. 2017, Azam, Haque et al. 2021, Lopez-Otin, Blasco et al. 2023). Age-related damaged mitochondria have been shown to affect mitochondrial energy metabolism and dynamics as well as redox homeostasis (Grimm and Eckert 2017). Brain aging research tools include diverse organisms models (Taormina, Ferrante et al. 2019). As mouse models have some limitations in terms of translatability to human physiology and lifespan, advanced human *in vitro* models are needed (Rydell-Tormanen and Johnson 2019). The inaccessibility of live human brain samples, such as neurons, has slowed progress in this area. Induced pluripotent stem cells (iPSCs) represent synthetic stem cells that are “reprogrammed” or induced from somatic cells cultured with various transcription factors like Klf4,c-Myc, Oct4 and Sox2 (Alciati, Reggiani et al. 2022). The first human iPSCs were created from a skin biopsy of skin fibroblasts (Abdullah, Pollock et al. 2012). Moreover, a number of biological correlates indicates that fibroblasts is an appropriate model for the study of the aging at the cellular level. The recent emergence of technologies that enable fibroblasts on one hand to be reprogrammed into iPSCs for neural cell generation and on the other hand to be converted directly into neural cells (induced neurons, iNs), offered a unique occasion to study crucial aspects of central nervous system function *in vitro* (Alciati, Reggiani et al. 2022). Mertens and colleagues showed that directly programmed iNs generated from aged donor fibroblasts retain at least donor population age-related transcriptional signatures and show functional impairments in their ability to properly compartmentalise nuclear and cytoplasmic proteins (Mertens, Paquola et al. 2015, Mertens, Marchetto et al. 2016), suggesting that direct conversion allows to identify the age-associated processes and the modelling of human aging at the cellular level. In contrast, it has been shown that the production of iPSCs reverses age-associated molecular features, such as telomere lengthening as well as age-related changes in the transcriptome (Puri and Wagner 2023).

However, conflicting data has been reported on the ability of conversion to reverse age-related modifications such as mitochondrial dysfunction, and to completely rejuvenate an aged somatic cell (Suhr, Chang et al. 2009). Thus, iPSCs might have the potential to carry epigenetic memory characteristic of the tissues from which they originate, which can affect their ability to differentiate (Puri and Wagner 2023).

Therefore, we wanted to investigate whether iPSCs from aged donors still exhibit their aging signatures at the mitochondrial level indicating a partial rejuvenation. Thus, we examined the mitochondrial bioenergetics and morphology in human iPSCs from elderly donors in comparison to those from young donors. We demonstrated that compared to human iPSCs from young donors, iPSCs from aged donors exhibit deficits in adenosine triphosphate (ATP) synthesis, a depolarisation of the mitochondrial membrane potential (MMP), exhibit increased generation in mitochondrial reactive oxygen species (ROS), and impairments in mitochondrial respiration and glycolytic remodelling. Moreover, we showed age-related modifications of mitochondrial network morphology, leading to more fragmented and shorter mitochondria.

## 2. Materials and methods

### 2.1 Chemicals and Reagents

Perkin Elmer (Waltham, MA, USA) supplied the ATPlite 1-step luminescence assay. DUTSCHER (Bernolsheim, France) supplied phosphate-buffered saline (PBS). Invitrogen (Waltham, MA, USA) supplied MitoSOX™ Red Mitochondrial Superoxide Indicator. Gibco (Waltham, MA, USA) provided the TrypLE™ Select Enzyme. Tetramethylrhodamine methyl ester perchlorate (TMRM), Hanks’ Balanced Salt Solution (HBSS), dihydrorhodamine 123 (DHR) were all were purchased from Sigma-Aldrich (St. Louis, MO, USA). Takara Bio (Kusatsu, Shiga, Japan) supplied the Cellartis DEF-CS 500 COAT-1 and the DEF-CS 500 culture system. Seahorse XF DMEM Medium, pH 7.4, Seahorse XF calibrant, Mito stress test kit, glutamine, glucose and pyruvate were supplied from Agilent (Santa Clara, CA, USA).

### 2.2 Human iPSC cells culture model

Takara, StemBANCC consortium and Dr Zameel Cader from the Oxford’s university kindly provided the iPSC lines (**Table 1**). Feeder-free conditions were used to maintained iPSCs. Plates were coated with DEF-CS COAT-1 and cells were grown using DEF-CS basal medium supplemented with GF-1 (1:333) and GF-2 (1:1000). To reduce the cell death after passing or plating, the medium was supplemented with the supplement GF-3 (1:1000). The cells were passaged once a week with TripLE Select. The culture medium was changed daily. A humidified incubator at 37 °C and 5 % CO2 was used for the incubation of the cells.

**Table 1.**
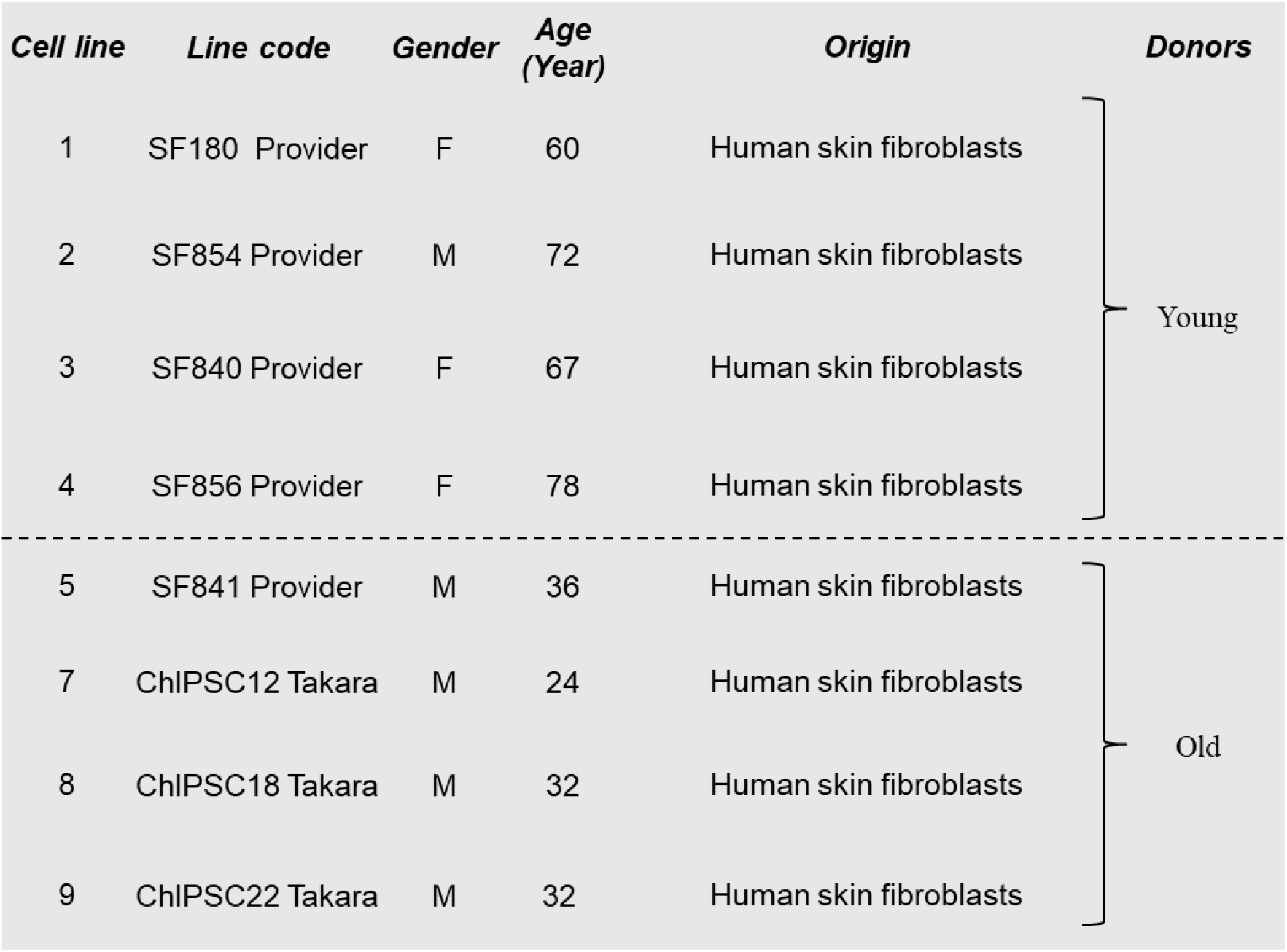
IPSC cell lines used in this study.

### 2.3 ATP Levels

A bioluminescence assay was used to determine the total ATP content of iPSC cells (Fairley, Lejri et al. 2023). Bioluminescence represents the measurement of the production of light by luciferase from ATP and luciferin. The Cytation 3 Cell Imaging Multi-mode Plate Reader (BioTek) was used to measure the emitted light, which was linearly related to the ATP concentration.The iPSCs from young and aged donors were plated each at a density of 20,000 cells/ well into coated white 96-well cell culture plate coated with COAT-1.

### 2.4 Determination of mitochondrial membrane potential (MMP)

Since the transmembrane distribution of MMP is dependent on MMP, it was measured using the potentiometric fluorescent TMRM. Cells were plated into a black 96-well cell culture plate coated with COAT-1 at a density of 20,000 cells/ well. The next day iPSC cells were incubate with TMRM (0.4 μM for 20 minutes). Fluorescence was measured using the multiplate reader Cytation 3 (BioTek) at 531 nm (excitation) / 595 nm (emission) after two washes with HBSS. The fluorescence intensity of the TMRM was dependent on the MMP.

### 2.5 Detection of reactive oxygen species levels (ROS)

Fluorescent dyes DHR and the Red Mitochondrial Superoxide Indicator (MitoSOX), were used to measure Levels of mitochondrial ROS, and mitochondrial superoxide anion radicals, respectively. IPSC cells were seeded at a density of 20000 cells/ well into black 96-well cell culture plates COAT-1 coated. At room temperature, the following day, iPSC cells were incubate with DHR (10 μM for 15 minutes) or MitoSOX (5 μM for 2 hours) in the dark on a shaker. After two washes with HBSS, the green fluorescent product generated by the oxidation of DHR was detected at 485 nm (excitation) / 538 nm (emission) using the Cytation 3 Cell Imaging Multi-mode Plate Reader (BioTek). At 531 nm (excitation) / 595 nm (emission), MitoSOX shows a red fluorescence. Fluorescence intensity was proportional to mitochondrial ROS levels and superoxide anion levels.

### 2.6 Bioenergetic phenotype

The Seahorse XF HS Mini Analyser (Agilent) was used to investigate key parameters related to mitochondrial respiration and glycolytic function. The oxygen consumption rate (OCR, mitochondrial respiration) was first measured in real-time separately from the extracellular acidification rate (ECAR, glycolysis). For both assays, cells (20,000 cells/well) were plated on Seahorse COAT-1 coated miniplates. The following day, for the OCR measurement, the XF Mito Stress Test protocol was carried out for the OCR measurement. The assay medium to measure the OCR the OCR consisted of the Seahorse XF DMEM medium at pH 7.4 and 18 mM glucose, 2 mM L-glutamine and 4 mM pyruvate. The glycolysis stress assay was performed for the ECAR measurement. The assay medium used to measure the ECAR was composed was composed of the Seahorse XF DMEM medium, with a pH of 7.4 and supplemented with 2 mM L-glutamine and 4 mM pyruvate. The OCR and ECAR were measured separately under basal conditions. To investigate the glycolytic function, sequential compound injections were performed to measure glycolysis, glycolytic capacity, and calculate of glycolytic reserve and non-glycolytic acidification. Bioenergetic parameters, especially the basal respiration, were automatically calculated on the Agilent Seahorse Analytics website.

### 2.7 Evaluation of mitochondrial membrane mass

The day before, iPSC cells (20 000 cells/well) were seeded in a 96 well plate COAT-1 coated. Mitochondrial membrane mass was determined using the dye MitoTracker Green FM (Molecular Probes, Invitrogen, Switzerland) (100 nM, 1 h). Fluorescence was measured using the Cytation 3 Cell Imaging Multi-mode Plate Reader (BioTek) at 490 nm (excitation) /516 nm (emission).

### 2.8 Mitochondrial Network Morphology

IPSCs were stained with the MitoTracker™ Red CMXRos to visualise and quantify the influence of aging on mitochondrial morphology. IPSCs from young and aged donors (20 000 cells/well) were plated COAT-1-coated 96-well cell culture plates. After 24h, 2% formaldehyde was used for cell fixation (for 15 minutes at room temperature). Then, permeabilisation was conducted (for 15 minutes at room temperature). The blocking process has been carried out for 1 hour at room temperature. For mitochondrial staining, cells were incubated with the the MitoTracker™ Red CMXRos in blocking buffer, for 30 minutes at room temperature. Nikon Eclipse Ti2-E inverted microscope with an oil objective was used to take images. Magnification was 94.5x (63x and 1.5x). FIJI software was used to assess changes in mitochondrial shape as previously described (Merrill, Flippo et al. 2017). Briefly, background subtraction has been applied to raw images (rolling ball radius = 10 pixels) and uneven mitochondrial labelling was ameliorated by local contrast enhancement using Contrast Limited Adaptive Histogram Equalisation (“CLAHE”). The “Tubeness” filter was applied for the segmentation of mitochondria. Then, the perimeter mitochondria and area were determined with the use of automatic thresholding and the “Analyze Particles” command. The “Skeletonize” function was applied to evaluate the mitochondrial length.

### 2.9 Statistical Analysis

The mean value ± SEM represents the data. Graph Pad Prism 9 was used for statistical analysis. The values have been normalised to the mean value of the iPSC from young donors group (=100%). Statistical comparisons were made using Student’s unpaired *t-test* between the iPSCs from young donors and the iPSCs from aged donors. Values are considered statistically significant if p-values < 0.05.

## 2. Results

### 2.9 IPSCs from aged donors display mitochondrial bioenergetic deficits

In iPSCs from young and old donors, ATP production (ATP), mitochondrial membrane potential (MMP) and reactive oxygen species (ROS) levels were evaluated for bioenergetics characterisation.

In the iPSCs from old donors, ATP production and MMP levels were significantly reduced by −36.8% and −36.25% respectively in comparison with the levels of iPSCs from young donors. (**Figure 1**). The fluorescence levels of DHR and MITOSOX were significantly elevated in old donor iPSCs compared to young donor iPSCs (**Figure 1**).

**Figure 1.**
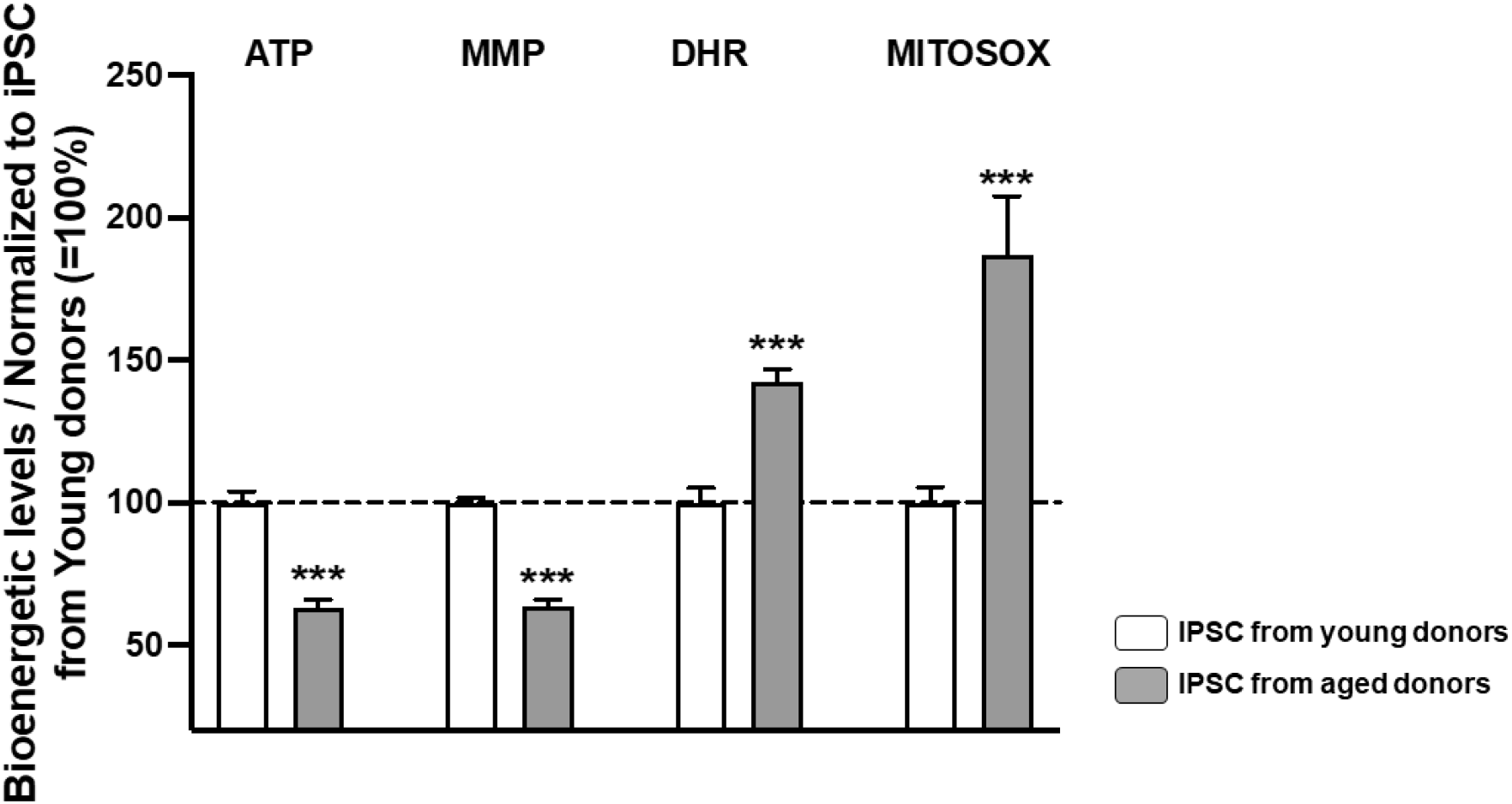
Bioenergetic readouts in iPSCs from young versus old donors including ATP generation (ATP), mitochondrial membrane potential (MMP), mitochondrial ROS (DHR) and mitochondrial superoxide anion (MITOSOX) levels. The values have been normalized to the iPSCs derived from young donors for each assay (=100%) and represented the mean ± SEM of three independent tests. Student *t-test* iPSCs from young donors vs iPSCs from aged donors ***P<0.001.

### 2.10 Decrease in mitochondrial respiration and improvement in glycolytic function in iPSCs from aged compared to young donors

Two main pathways, the mitochondrial oxidative phosphorylation (OXPHOS) and the cellular glycolysis, are needed to generate the ATP molecules for the iPSCs to ensure their energy demand. OCR measures mitochondrial respiration, ECAR glycolysis, and both were monitored separately in real time. We observed a significant reduction in the OCR (**Figure 2A**) of iPSCs from aged donors compared to iPSCs from young donors under basal conditions. Concerning the respiratory parameters (**Figure 2B**), iPSCs from aged donors demonstrated a significant decrease in basal respiration, spare respiratory capacity, maximal respiration, proton leak, ATP-production coupled respiration and non-mitochondrial oxygen consumption, when compared to iPSCs from young donors indicating a striking mitochondrial metabolic dysfunction.

**Figure 2.**
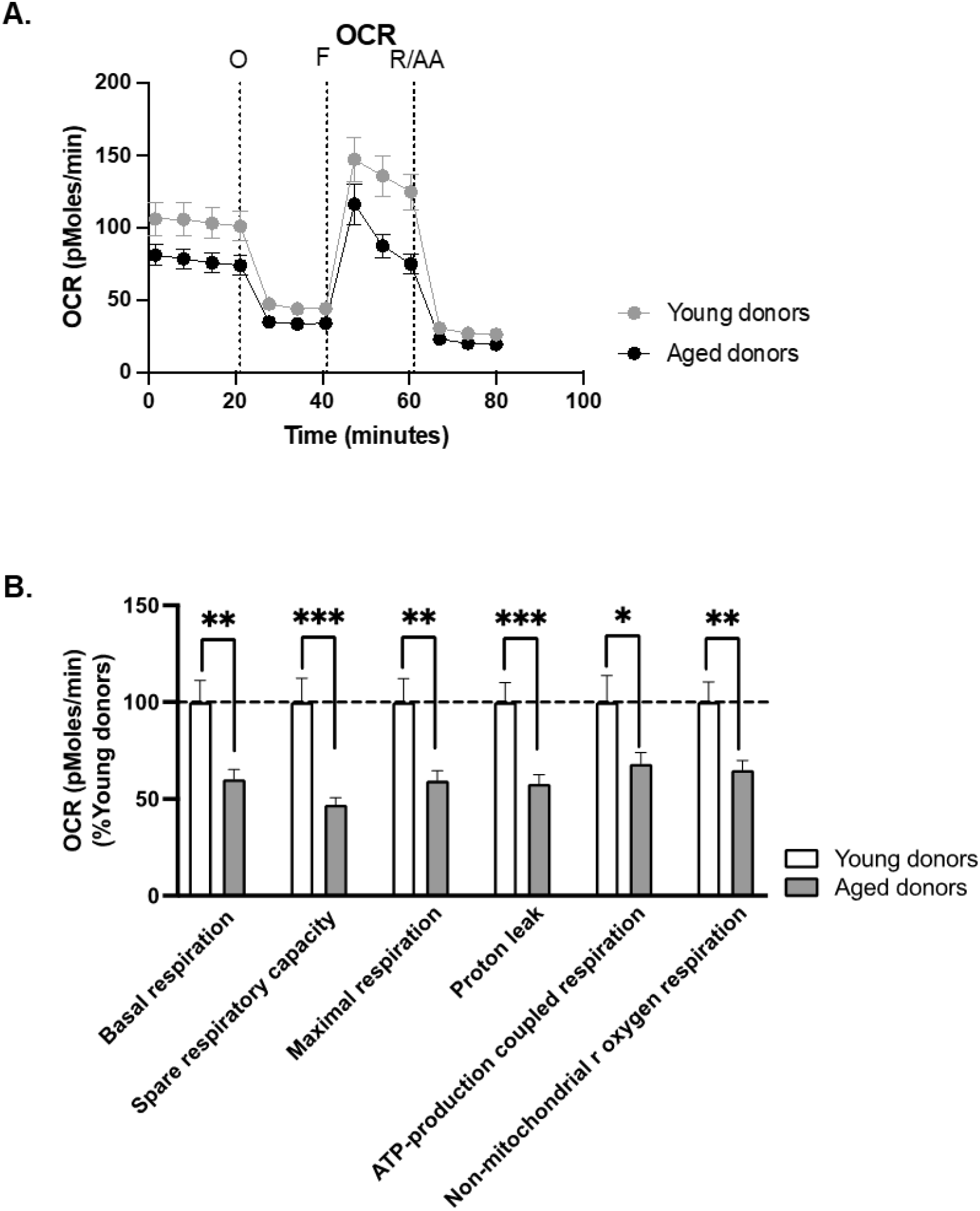
Analysis of seahorse data of (A) oxygen consumption rate (OCR), and (B) bioenergetic parameters of iPSCs from young and aged donors including basal respiration, spare respiratory capacity, maximal respiration, proton leak, ATP-production coupled respiration and non-mitochondrial oxygen consumption. To measure OCR in iPSCs from young and elderly donors, Mito Stress Test protocol was carried out. A decrease in the mitochondrial respiration and the respiratory parameters were observed in iPSCs from aged donors. Student *t-test* iPSCs from young donors vs iPSCs from aged donors *P<0.05, **P<0.01, ***P<0.001. OCR, Oxygen Consumption Rate, O: oligomycin, F: FCCP, R: rotenone, AA: antimycin A. Values represent the mean± SEM of three independent tests.

Interestingly, iPSCs from older donors compared to young donors showed high levels of ECAR. (**Figure 3A**), switching the IPSCs from aged donors to a more glycolytic state (**Figure 3B**).

**Figure 3.**
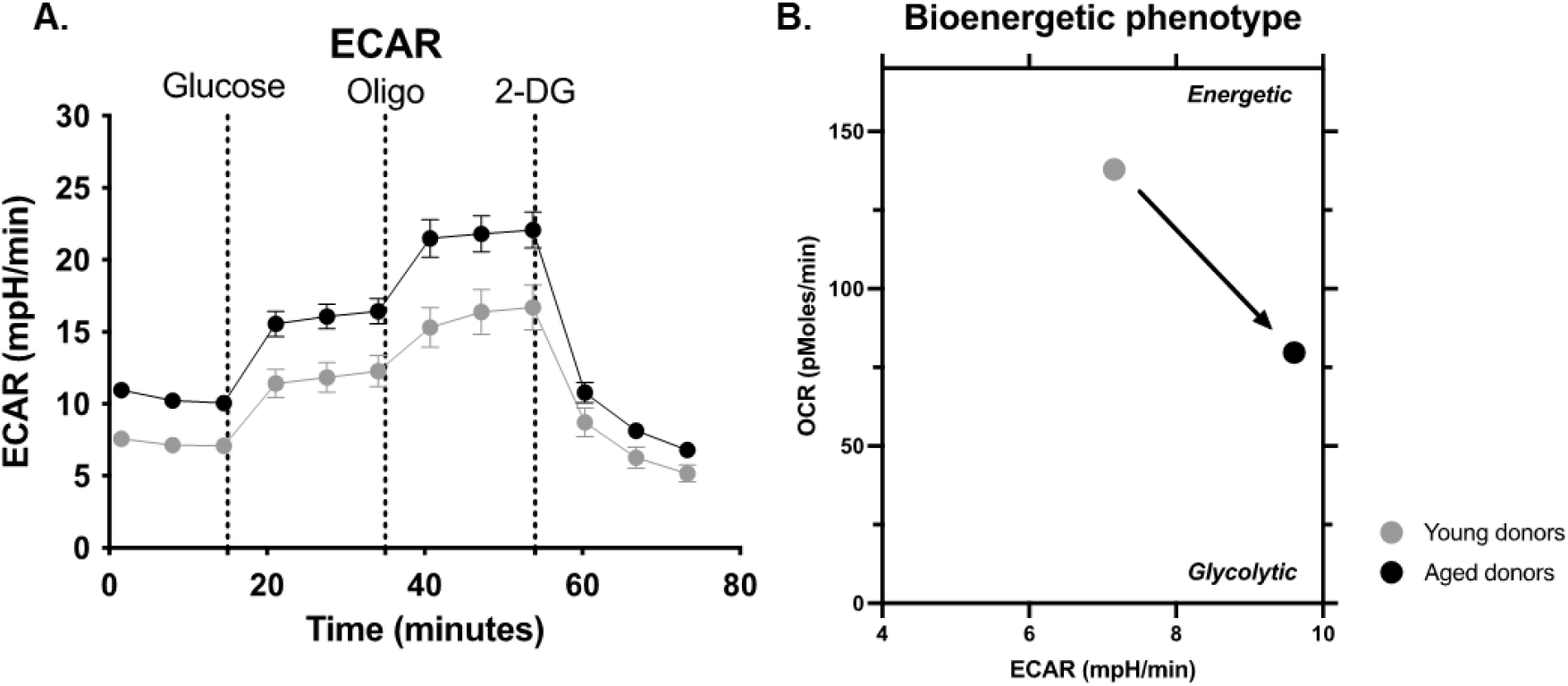
Seahorse analysis of (A) extracellular acidification rate (ECAR), (B) representative bioenergetics phenotype (mean value of the ECAR on the abscissa/mean value of the OCR (values from figure 2A) on the ordinate) in human iPSCs from young and aged donors. Seahorse Glycolysis Stress Test was performed to measure ECAR in iPSCs from young and aged donors following a sequential addition of rotenone/ antimycin A, and 2-deoxyglucose (2-DG). An increase in glycolytic function was observed in iPSCs from aged donors. ECAR, Extracellular Acidification Rate (Glycolysis), oligo: oligomycin. Values represent the mean± SEM of three independent experiments.

Together, these data suggest that the iPSCs from aged donors exerted a drop in the mitochondrial respiration and a “compensation” by increasing the glycolysis compared to the iPSCs from young donors to support the bioenergetic demands of the cell.

### 2.11 Preservation in mitochondrial mass in iPSCs from aged compared to young donors

Because these findings might be related to a decrease in mitochondrial mass in iPSCs from aged donors, we stained the mitochondria with the MitoTracker Green FM dye. We observed deacrased mitotracker staining in iPSCs from elderly donors compared to iPSCs from young donors, indicative of lower mitochondrial content in iPSCs from aged donors (**Figure 4B**). In fact, the iPSCs from aged donors showed a decrease in the mitochondrial mass with a reduction of −13% compared to the iPSCs from young donors (**Figure 4B**). Moreover, from the staining we received a first hint that the mitochondria appeared to be more fragmented in older donor iPSCs compared to young donor iPSCs (**Figure 4A**).

**Figure 4.**
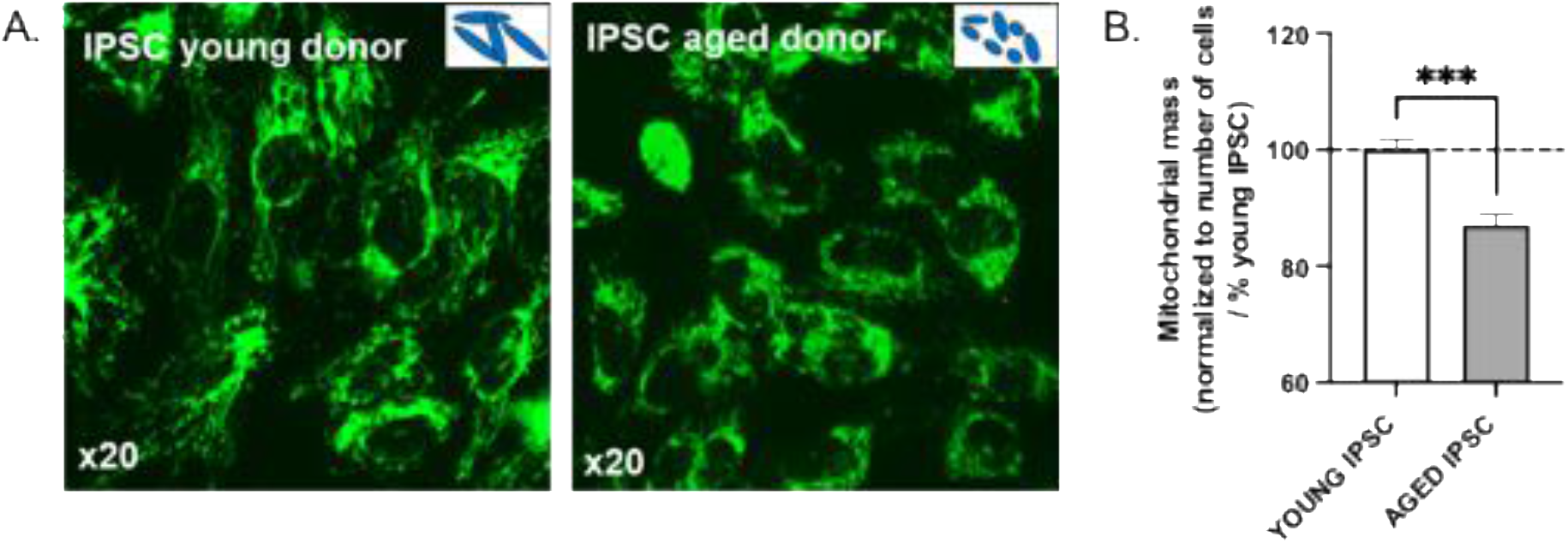
Characterization of the mitochondrial mass in iPSCs from young versus aged donors by using the MitoTracker Green FM. A significant reduction in mitochondrial mass was in iPSCs from older donors in comparison to those from young donors. Values represent the mean± SEM of three independent tests. Student *t-test* iPSCs from young donors vs iPSCs from aged donors ***P<0.001.

### 2.12 Age-related modification of the mitochondrial network morphology

Changes in the bioenergetic phenotype are closely linked to alterations in the mitochondrial morphology (Benard, Bellance et al. 2007). Lower bioenergetic status is often correlated with fragmented mitochondria and higher bioenergetic condition is associated with more elongated mitochondria (Galloway, Lee et al. 2012, Liu, McIntyre et al. 2020). Since iPSCs from aged donors showed bioenergetic deficiencies, we then evaluated the impact of aging on the morphology of the mitochondrial network using fluorescence microscopy. Mitochondrial marker staining was performed to visualise mitochondria by using CMXROS mitotracker red. Images were analysed to quantify the shape of the mitochondria through use of the FIJI software (**Figure 5A**).

**Figure 5.**
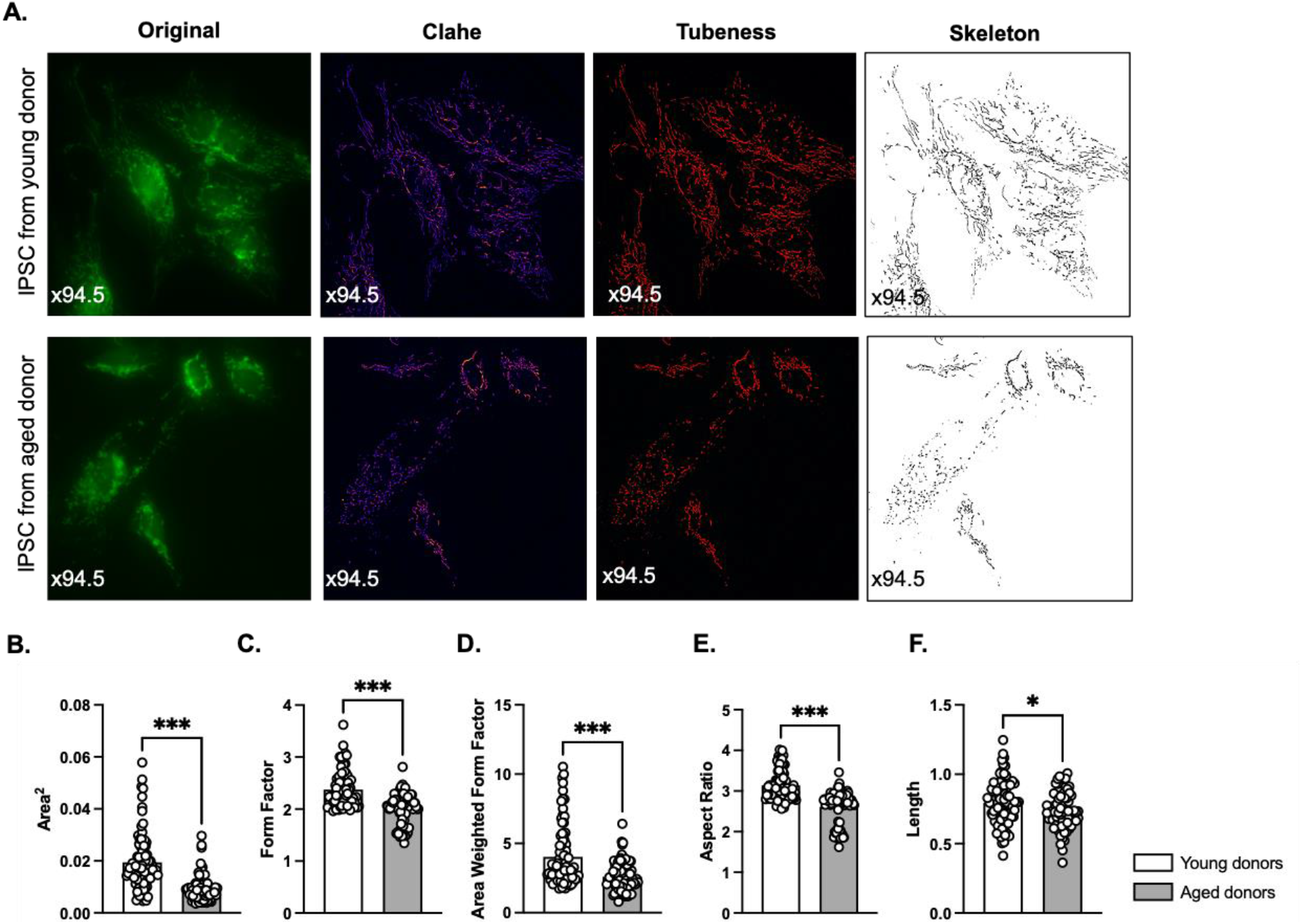
Impact of age on the morphology of the mitochondrial network in iPSCs. (A) Representative z-projection microscopy images of the stained mitochondrial network with the CMXROS mitotracker red (visualized in green by using FIJI) in iPSCs from young and aged donors (63× 1.5= 94.5 magnification). The original panels display mitochondrial network composite images (green). After processing with FIJI’s morphometry macro, the tubeness panels present the network of mitochondria (red). The skeleton panels display the mitochondrial network (grey) following further image processing in FIJI with Skeletonize function. (B–F) Quantification of mitochondrial network morphology parameters by assessment of (B) area^2^ (mean of the size of mitochondria), (C) form factor (elongation of mitochondria), (D) area-weighted form factor (a change in form factor towards larger mitochondrial size), (E) aspect ratio (Major/minor axis ratio), and (F) length (of mitochondria) in iPSCs from young and aged donors. Student *t-test* iPSCs from young donors vs iPSCs from aged donors*P<0.05, ***P<0.001.

All the parameters of the mitochondrial shape were significantly reduced, including the area^2^ (**Figure 5B**), the form factor (**Figure 5C**), the area-weighted form factor (**Figure 5D**), and the aspect ratio (**Figure 5E**) in iPSCs from aged donors compared to iPSCs from young donors. Additionally, we demonstrated a significant decrease in mitochondrial length (**Figure 5F**) in iPSCs from older donors in comparison to those from young donors. On the one hand, iPSCs from aged donors exhibited more fragmented mitochondria (**Figure 5A**), as indicated by a lower “form factor” value (**Figure 5C**) and a shorter length (**Figure 5F**) compared to iPSCs from young donors. On the other hand, the mitochondria of the iPSCs from young donors were more tubular in shape with higher form factor value (**Figure 5C**) and longer mitochondria (**Figure 5F**).

Overall, our data showed an age-related impact on mitochondrial dynamics and mitochondrial network morphology in iPSCs. Thereby, mitochondria are less elongated and in a more fragmented state.

## 3. Discussion

Many studies have focused mainly on age-related dysfunction in fibroblasts and directly derived neurons, but the mitochondrial impairment characterisation in human iPSCs from aged donors has been underexplored. (Mertens, Paquola et al. 2015, Mertens, Reid et al. 2018, Mertens, Herdy et al. 2021). Although it is proven that reprogramming towards cellular clock aging reset through pluripotency aging and removal of most of the cellular hallmarks associated with aging, a partial representation might be possible. Therefore, the aim was to evaluate whether iPSCs from aged donors still show their aging signature at the mitochondrial level. We characterized the age-related mitochondrial deficits in IPSCs from young and elderly donors on the bioenergetics, the oxidative stress and the mitochondrial mass and mitochondrial dynamics and network morphology. Our key findings were that compared to the iPSCs from young donors, iPSCs from elderly donors exhibited (1) lower ATP and MMP levels, an increase in the ROS production, (2) a low rate of mitochondrial respiration, and metabolic remodelling through high glycolytic functions as well as (3) impairments in the mitochondrial mass and mitochondrial morphology.

As cells age, the activity of the respiratory chain seems to reduce, thus increasing electron leakage and decreasing ATP production (Green, Galluzzi et al. 2011). The cell produces less energy, and at the same time, levels of oxidative stress rise, causing damage to other cellular metabolites (Grimm and Eckert 2017, Akbergenov, Duscha et al. 2018). Our results are consistent with data from ***in vitro studies models***. Masotti and colleagues investigated the ‘biological aging’ of iPSCs, specifically skin fibroblasts of a healthy individual, maintained *in vitro* for more than one year to represent “the aged” iPSCs and one month for the “young” iPSCs. They showed a decrease of the mitochondrial membrane potential levels in those “aged” iPSCs (Masotti, Celluzzi et al. 2014). Moreover, Petrini and colleagues suggested that “aged” iPSCs cultivated more than one year as a new model for the study of “premature” aging by evaluating the organization and expression pattern of nuclear envelope major constituents. However investigations at the mitochondrial level were missing (Petrini, Borghi et al. 2017). In our study, human iPSCs from aged donors exhibit ***in vivo aging phenotype*** such as lower ATP and MMP levels, whereas they exerted an augmentation of mitochondrial ROS and mitochondrial superoxide anion radicals levels in comparison with young donor human iPSCs. Interestingly, low level of ROS is implicated in the maintenance of the quiescent state of haematopoietic stem cells (HSCs), while increases in ROS serve as signalling molecules to support stem cells differentiation (Ushio-Fukai 2012). Excessive levels of ROS can create an oxidative microenvironment in pathophysiological states such as aging leading cellular damages (Ushio-Fukai 2012). Because of the high-energy demand of the iPSCs to maintain the cellular function, we next investigated the mitochondrial respiration and glycolysis in human iPSCs from young and aged donors. Rejuvenation allows aged cells to exert youthful characteristics through cellular reprogramming (Ji, Xiong et al. 2023). Our results have shown that iPSCs derived from older donors exerted low mitochondrial respiration compared to iPSCs from young donors. Notably, our study is the first to assess the glycolytic parameter in human iPSCs from aged donors highlighting a partial rejuvenation through glycolytic remodelling in these cells compared to the human iPSCs from young donors. Indeed, we were able to show that human iPSCs from old donors used increased glycolysis as a compensatory mechanism for the reduction in OXPHOS (oxidative phosphorylation) to respond to high energy demands compared to those from young donors. In line with our data, iPSCs are proliferative cells and ensure self-renewal and pluripotency processes through the aerobic glycolysis for the production of energy with low levels of OXPHOS (Ishida, Nakao et al. 2020). Moreover, metabolic modification from OXPHOS to glycolysis in iPSCs can be explained by the Warburg effect which is essential for the maintenance of the stem cell characteristics (Prigione, Fauler et al. 2010, Varum, Rodrigues et al. 2011, Panopoulos, Yanes et al. 2012, Ishida, Nakao et al. 2020).

Further research is now required to complete these findings, first on the corresponding fibroblasts and iPSCs-derived neurons from the same human young and aged donors in order to do comparative analysis to explore the preservation of aging fealures more deeply. The advantage of using iPSCs is that they can be maintained indefinitely with a high cell number and stored easily compared to the iNs which are known to be limited in the cell number, not suitable for screening. iPSCs have the ability to differentiate into neural progenitor cells (NPC), also a potential model that can be a useful tool for the screening drugs.

Taking together, our data revealed that iPSCs an aged donors could still retain age-associated changes in mitochondrial function, thus displaying the aging signature in the absence of a full rejuvenation. Notably, aged iPSCs exhibited impairments in mitochondrial bioenergetics, resulting in a mitochondrial respiratory failure with concomitant ATP depletion, which was accompanied by a depolarization of the MMP and an augmented production of different mitochondrial ROS as well as mitochondrial network alterations. Overall, the preservation of the mitochondrial aging signature in iPSCs from aged donors offers opportunities for drug screening to promote healthy aging and in the long term combat against age-related diseases at the mitochondrial levels.

## ACKNOWLEDGMENTS

We kindly thank Mrs Fides Meier for general laboratory coordination.

## AUTHOR CONTRIBUTIONS

Conceptualization: A.E.; Methodology: I.L.; Investigation: I.L.; Formal analysis: I.L.; Writing—original draft: I.L.; Funding acquisition: A.E.; Resources: A.E.; Supervision: A.E. All authors have read and agreed to the published version of the manuscript.

## INSTITUTIONAL REVIEW BOARD STATEMENT

Not applicable.

## INFORMED CONSENT STATEMENT

Not applicable.

## DATA AVAILABILITY STATEMENT

The data presented in this study are available on request from the corresponding author.

## References

Abdullah, A. I., A. Pollock and T. Sun (2012). “The path from skin to brain: generation of functional neurons from fibroblasts.” Mol Neurobiol 45(3): 586–595.

Alciati, A., A. Reggiani, D. Caldirola and G. Perna (2022). “Human-Induced Pluripotent Stem Cell Technology: Toward the Future of Personalized Psychiatry.” J Pers Med 12(8).

Azam, S., M. E. Haque, R. Balakrishnan, I. S. Kim and D. K. Choi (2021). “The Ageing Brain: Molecular and Cellular Basis of Neurodegeneration.” Front Cell Dev Biol 9: 683459.

Benard, G., N. Bellance, D. James, P. Parrone, H. Fernandez, T. Letellier and R. Rossignol (2007). “Mitochondrial bioenergetics and structural network organization.” J Cell Sci 120(Pt 5): 838–848.

Fairley, L. H., I. Lejri, A. Grimm and A. Eckert (2023). “Spermidine Rescues Bioenergetic and Mitophagy Deficits Induced by Disease-Associated Tau Protein.” Int J Mol Sci 24(6).

Galloway, C. A., H. Lee and Y. Yoon (2012). “Mitochondrial morphology-emerging role in bioenergetics.” Free Radic Biol Med 53(12): 2218–2228.

Grimm, A. and A. Eckert (2017). “Brain aging and neurodegeneration: from a mitochondrial point of view.” J Neurochem 143(4): 418–431.

Ishida, T., S. Nakao, T. Ueyama, Y. Harada and T. Kawamura (2020). “Metabolic remodeling during somatic cell reprogramming to induced pluripotent stem cells: involvement of hypoxia-inducible factor 1.” Inflamm Regen 40: 8.

Ji, S., M. Xiong, H. Chen, Y. Liu, L. Zhou, Y. Hong, M. Wang, C. Wang, X. Fu and X. Sun (2023). “Cellular rejuvenation: molecular mechanisms and potential therapeutic interventions for diseases.” Signal Transduct Target Ther 8(1): 116.

Kauppila, T. E. S., J. H. K. Kauppila and N. G. Larsson (2017). “Mammalian Mitochondria and Aging: An Update.” Cell Metab 25(1): 57–71.

Liu, Y. J., R. L. McIntyre, G. E. Janssens and R. H. Houtkooper (2020). “Mitochondrial fission and fusion: A dynamic role in aging and potential target for age-related disease.” Mech Ageing Dev 186: 111212.

Lopez-Otin, C., M. A. Blasco, L. Partridge, M. Serrano and G. Kroemer (2023). “Hallmarks of aging: An expanding universe.” Cell 186(2): 243–278.

Masotti, A., A. Celluzzi, S. Petrini, E. Bertini, G. Zanni and C. Compagnucci (2014). “Aged iPSCs display an uncommon mitochondrial appearance and fail to undergo in vitro neurogenesis.” Aging (Albany NY) 6(12): 1094–1108.

Merrill, R. A., K. H. Flippo and S. Strack (2017). Measuring Mitochondrial Shape with Image J.

Mertens, J., J. R. Herdy, L. Traxler, S. T. Schafer, J. C. M. Schlachetzki, L. Bohnke, D. A. Reid, H. Lee, D. Zangwill, D. P. Fernandes, R. K. Agarwal, R. Lucciola, L. Zhou-Yang, L. Karbacher, F. Edenhofer, S. Stern, S. Horvath, A. C. M. Paquola, C. K. Glass, S. H. Yuan, M. Ku, A. Szucs, L. S. B. Goldstein, D. Galasko and F. H. Gage (2021). “Age-dependent instability of mature neuronal fate in induced neurons from Alzheimer’s patients.” Cell Stem Cell 28(9): 1533–1548 e1536.

Mertens, J., M. C. Marchetto, C. Bardy and F. H. Gage (2016). “Evaluating cell reprogramming, differentiation and conversion technologies in neuroscience.” Nat Rev Neurosci 17(7): 424–437.

Mertens, J., A. C. M. Paquola, M. Ku, E. Hatch, L. Bohnke, S. Ladjevardi, S. McGrath, B. Campbell, H. Lee, J. R. Herdy, J. T. Goncalves, T. Toda, Y. Kim, J. Winkler, J. Yao, M. W. Hetzer and F. H. Gage (2015). “Directly Reprogrammed Human Neurons Retain Aging-Associated Transcriptomic Signatures and Reveal Age-Related Nucleocytoplasmic Defects.” Cell Stem Cell 17(6): 705–718.

Mertens, J., D. Reid, S. Lau, Y. Kim and F. H. Gage (2018). “Aging in a Dish: iPSC-Derived and Directly Induced Neurons for Studying Brain Aging and Age-Related Neurodegenerative Diseases.” Annu Rev Genet 52: 271–293.

Panopoulos, A. D., O. Yanes, S. Ruiz, Y. S. Kida, D. Diep, R. Tautenhahn, A. Herrerias, E. M. Batchelder, N. Plongthongkum, M. Lutz, W. T. Berggren, K. Zhang, R. M. Evans, G. Siuzdak and J. C. Izpisua Belmonte (2012). “The metabolome of induced pluripotent stem cells reveals metabolic changes occurring in somatic cell reprogramming.” Cell Res 22(1): 168–177.

Petrini, S., R. Borghi, V. D’Oria, F. Restaldi, S. Moreno, A. Novelli, E. Bertini and C. Compagnucci (2017). “Aged induced pluripotent stem cell (iPSCs) as a new cellular model for studying premature aging.” Aging (Albany NY) 9(5): 1453–1469.

Prigione, A., B. Fauler, R. Lurz, H. Lehrach and J. Adjaye (2010). “The senescence-related mitochondrial/oxidative stress pathway is repressed in human induced pluripotent stem cells.” Stem Cells 28(4): 721–733.

Puri, D. and W. Wagner (2023). “Epigenetic rejuvenation by partial reprogramming.” Bioessays 45(4): e2200208.

Rydell-Tormanen, K. and J. R. Johnson (2019). “The Applicability of Mouse Models to the Study of Human Disease.” Methods Mol Biol 1940: 3–22.

Suhr, S. T., E. A. Chang, R. M. Rodriguez, K. Wang, P. J. Ross, Z. Beyhan, S. Murthy and J. B. Cibelli (2009). “Telomere dynamics in human cells reprogrammed to pluripotency.” PLoS One 4(12): e8124.

Taormina, G., F. Ferrante, S. Vieni, N. Grassi, A. Russo and M. G. Mirisola (2019). “Longevity: Lesson from Model Organisms.” Genes (Basel) 10(7).

Ushio-Fukai, M. (2012). Reactive oxygen species and stem/progenitor cells.

Varum, S., A. S. Rodrigues, M. B. Moura, O. Momcilovic, C. A. t. Easley, J. Ramalho-Santos, B. Van Houten and G. Schatten (2011). “Energy metabolism in human pluripotent stem cells and their differentiated counterparts.” PLoS One 6(6): e20914.

